# The language network is recruited but not required for non-verbal event semantics

**DOI:** 10.1101/696484

**Authors:** Anna A. Ivanova, Zachary Mineroff, Vitor Zimmerer, Nancy Kanwisher, Rosemary Varley, Evelina Fedorenko

## Abstract

The ability to combine individual meanings into complex representations of the world is often associated with language. Yet people also construct combinatorial event-level representations from non-linguistic input, e.g. from visual scenes. Here, we test whether the language network in the human brain is involved in and necessary for semantic processing of nonverbal events. In Experiment 1, we scanned participants with fMRI while they performed a semantic plausibility judgment task vs. a difficult perceptual control task on sentences and line drawings that describe/depict simple agent-patient interactions. We found that the language network responded robustly during the semantic task but not during the perceptual control task. This effect was observed for both sentences and pictures (although the response to sentences was stronger). Thus, language regions in healthy adults are engaged during a semantic task performed on pictorial depictions of events. But is this engagement necessary? In Experiment 2, we tested two individuals with global aphasia, who have sustained massive damage to perisylvian language areas and display severe language difficulties, against a group of age-matched control participants. Individuals with aphasia were severely impaired on a task of matching sentences and pictures. However, they performed close to controls in assessing the plausibility of pictorial depictions of agent-patient interactions. Overall, our results indicate that the left fronto-temporal language network is recruited but not necessary for semantic processing of nonverbal events.

## Introduction

Many thinkers have argued for an intimate relationship between language and thought, in fields as diverse as philosophy (Carruthers, 2002; Davidson, 1975; Wittgenstein, 1961), psychology (Sokolov, 1972; Vygotski, 2012; Watson, 1920), linguistics (Berwick & Chomsky, 2016; Bickerton, 1990; Chomsky, 2007; Hinzen, 2013; Jackendoff, 1996), and artificial intelligence (Brown et al., 2020; Goldstein & Papert, 1977; Turing, 1950; Winograd, 1976). According to such accounts, language enables us to access our vast knowledge of objects, ideas, and relations — often referred to as semantic knowledge — and flexibly combine individual semantic units to produce complex thoughts. In fact, the process of accessing semantic knowledge is commonly equated with the process of extracting meanings from words, phrases, and sentences (Altshuler et al., 2019; Binder et al., 2009; Fillmore, 2006; Milberg & Blumstein, 1981; Pinker & Levin, 1991; Talmy, 2000). Here, we test the link between language and thought by examining the role of the language network in a non-verbal combinatorial semantic task.

Recent evidence from neuroscience suggests that language processing is largely distinct from other aspects of cognition (Fedorenko & Blank, 2020; Fedorenko & Varley, 2016). A network of frontal and temporal brain regions (here referred to as the ‘language network’) has been found to respond to written/spoken/signed words and sentences, but not to mental arithmetic, music perception, executive function tasks, action/gesture perception, or computer programming (Amalric & Dehaene, 2019; Fedorenko et al., 2011; Ivanova et al., 2020; Jouravlev et al., 2019; Liu et al., 2020; Monti et al., 2009, 2012; Pritchett et al., 2018). Similarly, investigations of patients with profound disruption of language capacity (global aphasia) have shown that these individuals can solve arithmetic and logic problems, appreciate music, and think about others’ thoughts in spite of their language impairments (Basso & Capitani, 1985; Luria et al., 1965; Varley et al., 2005; Varley & Siegal, 2000), providing converging evidence that language and thought are neurally distinct.

Despite this significant progress in dissociating linguistic and non-linguistic processing, the role of the language network in non-verbal semantics remains unclear. Neuroimaging studies that explicitly compared verbal and non-verbal semantic processing (Devereux et al., 2013; Fairhall & Caramazza, 2013; Handjaras et al., 2017; Vandenberghe et al., 1996; Visser et al., 2012, among others) often reported overlapping activation in left-lateralized frontal and temporal areas, which may reflect the engagement of the language network in non-verbal cognition. However, these studies have typically relied on group analyses — an approach known to overestimate overlap in cases of nearby functionally distinct areas (Nieto-Castañón & Fedorenko, 2012) — and/or do not report effect sizes, which are critical for interpreting the functional profiles of the regions in question (a region that responds similarly strongly to verbal and non-verbal semantic tasks plausibly supports computations that are different from a region that responds to both, but shows a 2-3 times stronger response to verbal semantics; see, e.g., Chen et al. (2017) for discussion). Furthermore, as in any neuroimaging study, these findings do not speak to the causal role of the language network in non-verbal semantic processing (cf. Weber & Thompson-Schill, 2010). Meanwhile, neuropsychology studies have often reported dissociations between linguistic and semantic deficits in patients with aphasia (e.g., Antonucci & Reilly, 2008; Jefferies & Lambon Ralph, 2006; Warrington, 1975), suggesting that linguistic and semantic processes rely on distinct neural circuits. However, given that language abilities are only partially impaired in most reported cases of aphasia, it is hard to completely rule out the involvement of the language network in semantic processing.

Further, very few neuroimaging (Baldassano et al., 2018; Thierry & Price, 2006) or neuropsychological (Dresang et al., 2019; Marshall et al., 1993) studies have investigated the relationship between linguistic and non-linguistic semantic processing of events (as opposed to individual objects or actions). Constructing event-level meaning representations requires object and action processing but is not reducible to them (Dresang et al., 2019), and therefore may engage additional mental operations. In particular, to understand an event, we must identify *relations* between participating entities and assign them *thematic roles* (Estes et al., 2011). This process of identifying who did what to whom has traditionally been considered a key function of the language system (Fillmore, 1968; Gruber, 1965; Jackendoff, 1990). Thus, if any aspect of semantic processing requires language, event understanding would seem to be one of the strongest candidates.

Event processing has perhaps been most extensively investigated in EEG research, where a number of studies have reported that semantic violations in visually presented scenes/events evoke the N400 response (e.g., Coco et al., 2020; Cohn, 2020; Proverbio & Riva, 2009; Võ & Wolfe, 2013; Sitnikova et al., 2008; West & Holcomb, 2002; see Kutas & Federmeier, 2011, for a review), similarly to semantic violations in sentences, where the N400 component was originally discovered (Kutas & Hillyard, 1980). The EEG results have been taken to suggest that linguistic and visual semantic processing rely on a shared mechanism. However, because the neural generators of the N400 remain debated (Lau et al., 2008, 2016; Matsumoto et al., 2005; Zhu et al., 2019), this evidence does not definitively demonstrate the involvement of the language network in visual event processing.

Here, we synergistically combine neuroimaging and neuropsychological evidence to ask whether the language network is engaged during and/or necessary for non-verbal event semantics. We focus on the understanding of agent-patient relations (“who did what to whom”) in visually presented scenes. Identification of thematic relations is critical to understanding and generating sentences (Carlson & Tanenhaus, 1988; Fillmore, 2002; Jackendoff, 1987), but “agent” and “patient” are not exclusively linguistic notions: they likely constitute part of humans’ core knowledge (Rissman & Majid, 2019; Spelke & Kinzler, 2007; Strickland, 2017; L. Wagner & Lakusta, 2009) and are integral to visual event processing (Cohn & Paczynski, 2013; Hafri et al., 2018). Investigating the role of the language network in processing agent-patient relations therefore constitutes a crucial test of the relationship between language and combinatorial event semantics.

We used two kinds of evidence in our study: (1) fMRI in neurotypical participants, and (2) behavioral evidence from two individuals with global aphasia and a group of age-matched healthy controls. All participants were asked to evaluate the plausibility of events, presented either as sentences (neurotypicals only) or pictures. Plausibility was manipulated by varying the typicality of agent and patient roles (e.g., a cop arresting a criminal vs. a criminal arresting a cop). To foreshadow our results, we find that language-responsive brain areas in neurotypical participants respond during the plausibility task for both sentences and pictures (although the responses are lower for pictures). However, individuals with global aphasia, who sustained severe damage to language areas, perform well on the picture plausibility task, suggesting that the language network is not required for extracting complex semantic information from visual depictions of events.

## Results

### Experiment 1: Is the language network active during semantic processing of events?

In the first experiment, we presented twenty-one neurotypical participants with sentences and pictures describing/depicting agent-patient interactions that were either plausible or implausible (**Figure 1**), while the participants were undergoing an fMRI scan. Participants performed a semantic judgment task on the sentences and pictures, as well as a difficulty-matched low-level perceptual control task on the same stimuli, in a 2×2 blocked design. In separate blocks, participants were instructed to indicate either i) whether the stimulus was plausible or implausible (the semantic task) or ii) whether the stimulus was moving slowly to the left or right (the perceptual task). The language network in each participant was identified using a separate functional language localizer task (sentences > nonwords contrast; Fedorenko et al., 2010). We then measured the response of those regions to sentences and pictures during the semantic and perceptual tasks.

**Figure 1.**
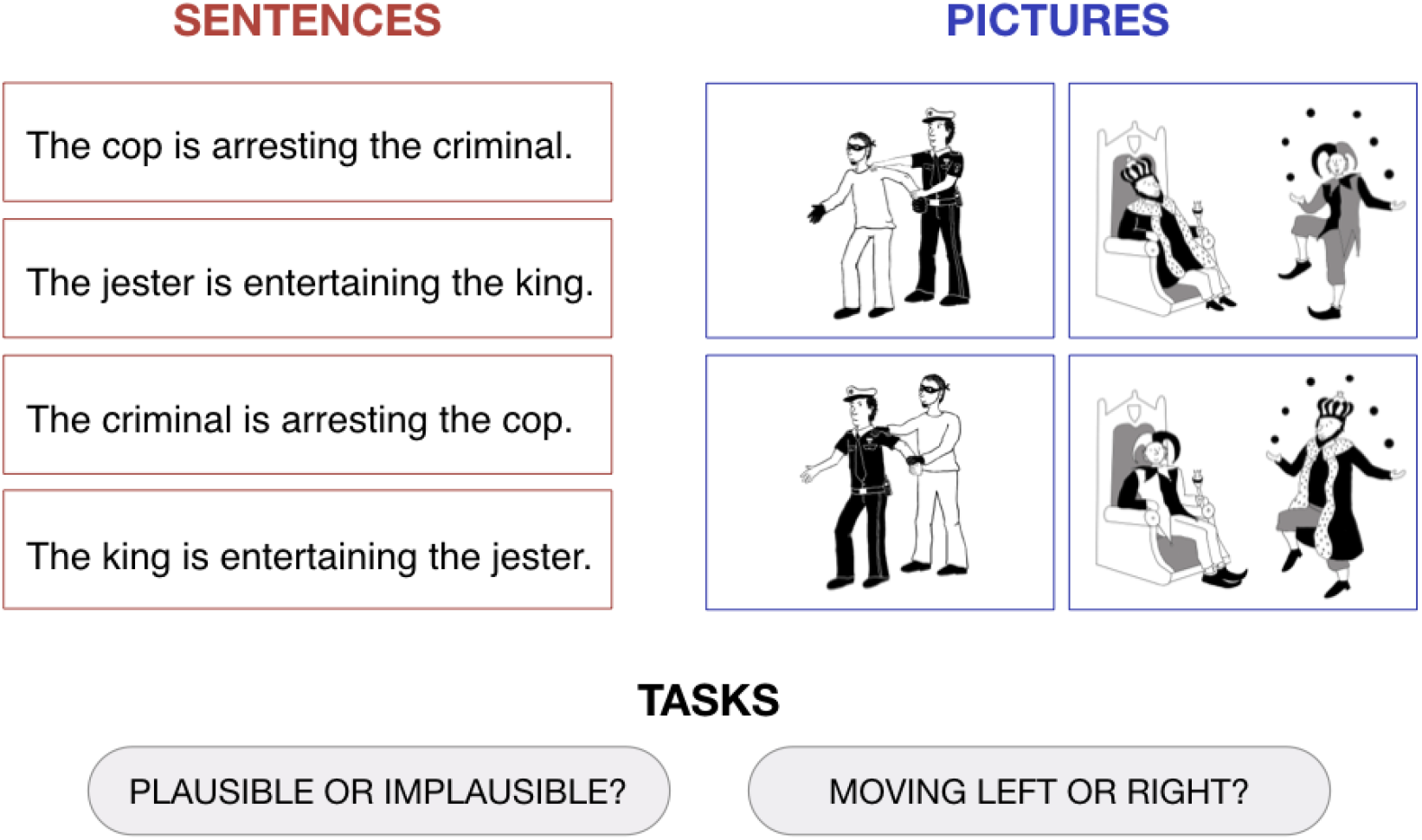
Sample stimuli used in the experiment. For both sentences and pictures, participants were required to perform either a semantic plausibility judgment task (“Plausible or implausible?”) or a control perceptual task (“Moving left or right?”). The full set of materials is available on the paper website (https://osf.io/gsudr/).

Although diverse non-linguistic tasks have been previously shown not to engage the language network (Fedorenko & Varley, 2016), we here found that the language regions responded more strongly during the semantic task on both sentences and pictures compared to the perceptual control task (**Figure 2A**). A linear mixed-effects model, with task and stimulus type as fixed effects and fROI and participant as random effect intercepts, showed a significant effect of task (semantic > perceptual, *β* = 0.71, *p* < .001). Responses to sentences during the semantic task were stronger than responses to pictures, as indicated by the interaction between task and stimulus type (*β* = .43, *p* = 0.025), although there was no main effect of stimulus type (sentences > pictures; *β* = .01, *p* = 0.925). The regression model intercept (pictures, perceptual task) was not significantly different from 0 (*β* = .28, *p* = 0.269), indicating that the language network was activated by the semantic task rather than deactivated by the perceptual task. These results demonstrate that, in the context of a semantic task, the language network responds not only to sentences, but also to pictures.

**Figure 2.**
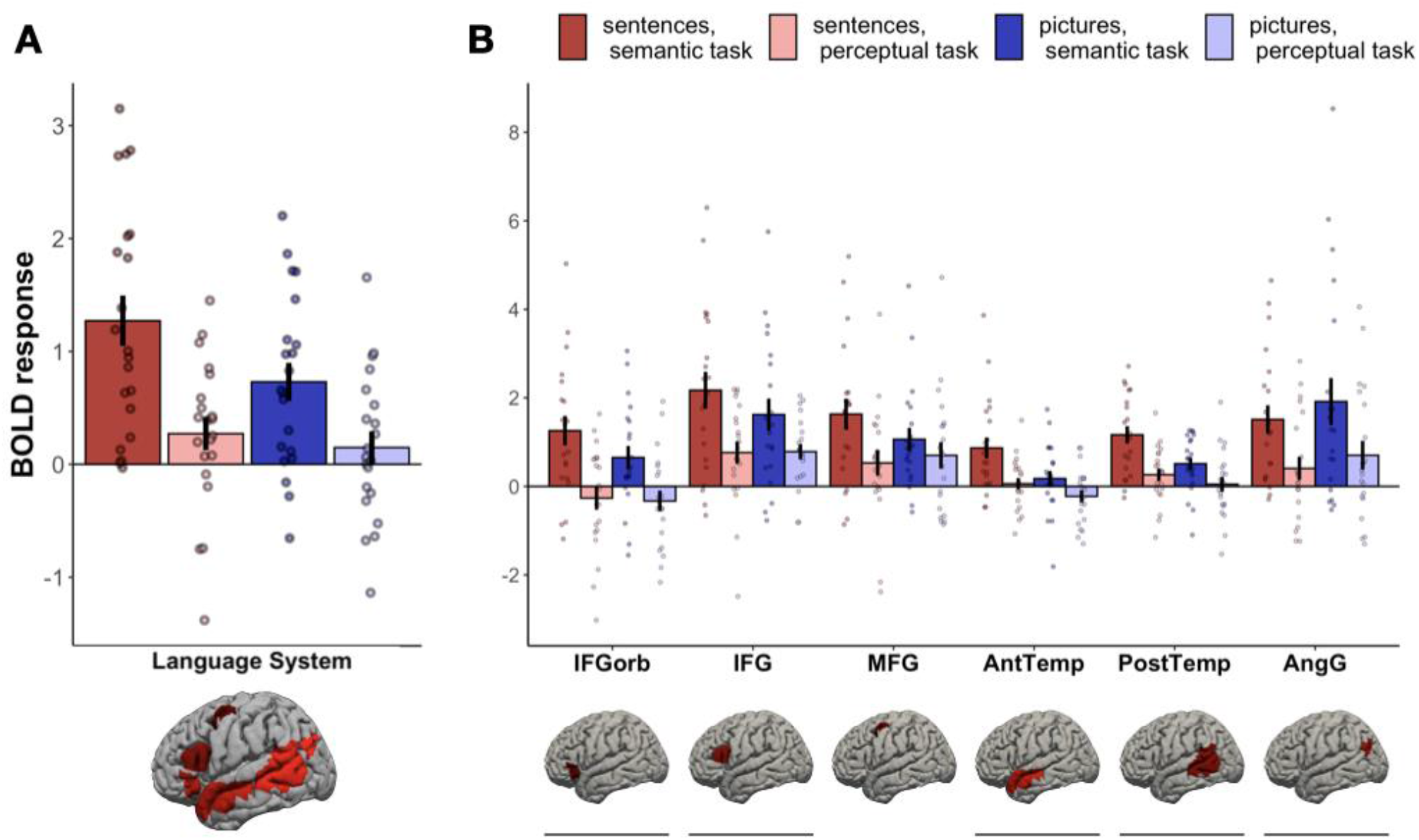
BOLD response during the four task conditions within (A) the language network as a whole and (B) each of the six language fROIs. The underlined fROIs show significantly higher activation for the semantic task than for the perceptual task, across both sentences and pictures. The fROI labels correspond to approximate anatomical locations: IFGorb – the orbital portion of the left inferior frontal gyrus; IFG – left inferior frontal gyrus; MFG – left middle frontal gyrus; AntTemp – left anterior temporal cortex; PostTemp – left posterior temporal cortex; AngG – left angular gyrus. Within each parcel, the responses to the critical experiment conditions are extracted from the top 10% most language-responsive voxels (selected in each of the 21 individuals separately). Error bars indicate standard error of the mean; dots indicate individual participants’ responses.

To investigate the different brain regions comprising the language network, we conducted follow-up analyses on individual fROIs’ activity (FDR-corrected for the number of regions) (**Figure 2B**). These revealed a significant semantic > perceptual task effect in all fROIs except the one in the middle frontal gyrus (**Table 1**). The effect of stimulus type and the interaction between task and stimulus type were not significant in any fROI, although, numerically, responses to sentences during the semantic task were stronger than responses to any other condition in all fROIs except the left angular gyrus.

**Table 1.** Regression model terms for fROI-based statistical analyses. The p-value is FDR-corrected for the number of regions. Significant terms are highlighted in bold. The fROI labels correspond to the approximate anatomical locations: IFGorb – the orbital portion of the left inferior frontal gyrus; IFG – left inferior frontal gyrus; MFG – left middle frontal gyrus; AntTemp – left anterior temporal cortex; PostTemp – left posterior temporal cortex; AngG – left angular gyrus.

| ROI | Regression Term | Beta | p-value |
| --- | --- | --- | --- |
| IFGorb | Intercept | -0.33 | 0.267 |
|  | Stimulus (Sent>Pic) | 0.07 | 0.949 |
|  | **Task (Sem>Perc)** | **0.98** | **0.01** |
|  | Stimulus:Task | 0.54 | 0.283 |
| IFG | Intercept | 0.78 | 0.067 |
|  | Stimulus (Sent>Pic) | -0.02 | 0.949 |
|  | **Task (Sem>Perc)** | **0.84** | **0.014** |
|  | Stimulus:Task | 0.57 | 0.28 |
| MFG | Intercept | 0.7 | 0.067 |
|  | Stimulus (Sent>Pic) | -0.17 | 0.739 |
|  | Task (Sem>Perc) | 0.36 | 0.147 |
|  | Stimulus:Task | 0.74 | 0.231 |
| AntTemp | Intercept | -0.23 | 0.267 |
|  | Stimulus (Sent>Pic) | 0.28 | 0.732 |
|  | **Task (Sem>Perc)** | **0.4** | **0.037** |
|  | Stimulus:Task | 0.41 | 0.24 |
| PostTemp | Intercept | 0.05 | 0.766 |
|  | Stimulus (Sent>Pic) | 0.21 | 0.739 |
|  | **Task (Sem>Perc)** | **0.46** | **0.026** |
|  | Stimulus:Task | 0.44 | 0.24 |
| AngG | Intercept | 0.7 | 0.122 |
|  | Stimulus (Sent>Pic) | -0.3 | 0.739 |
|  | **Task (Sem>Perc)** | **1.21** | **0.004** |
|  | Stimulus:Task | -0.11 | 0.823 |

We also performed a random effects whole-brain group analysis (**Figure S1**), which yielded results similar to the fROI-based analyses described above. Specifically, we found that the semantic > perceptual contrast for both sentences and pictures activates left-lateralized frontal and temporal regions that overlap with the language parcels (used to constrain the definition of individual language fROIs). The extent of semantics-evoked activation in the left lateral temporal areas was weaker for pictures than sentences (the opposite was true on the ventral surface of the left temporal lobe).

Overall, the first experiment revealed that the language network is strongly and significantly recruited for semantic processing of events presented not only verbally (through sentences), but also non-verbally (through pictures). Specifically, the language network is active when we access the meanings of pictures that depict agent-patient interactions and relate them to stored world knowledge about these protagonists. It is worth noting, however, that responses to the semantic task are stronger for sentences than for pictures (**Figure 2B**), suggesting that the language network may play a less important role in non-verbal semantic processing. To test whether the engagement of the language network is *necessary* for comprehending visually presented events, we turn to behavioral evidence from individuals with global aphasia.

### Experiment 2: Is the language network necessary for semantic processing of events?

In the second experiment, we examined two individuals with global aphasia, a disorder characterized by severe linguistic impairments. Both individuals (S.A. and P.R.) had suffered large vascular lesions that resulted in extensive damage to left perisylvian cortex, including the language network (see **Figure 3** for lesion images, including a probabilistic map of the language network based on fMRI data from neurotypical participants, overlayed onto one participant’s MRI).

**Figure 3.**
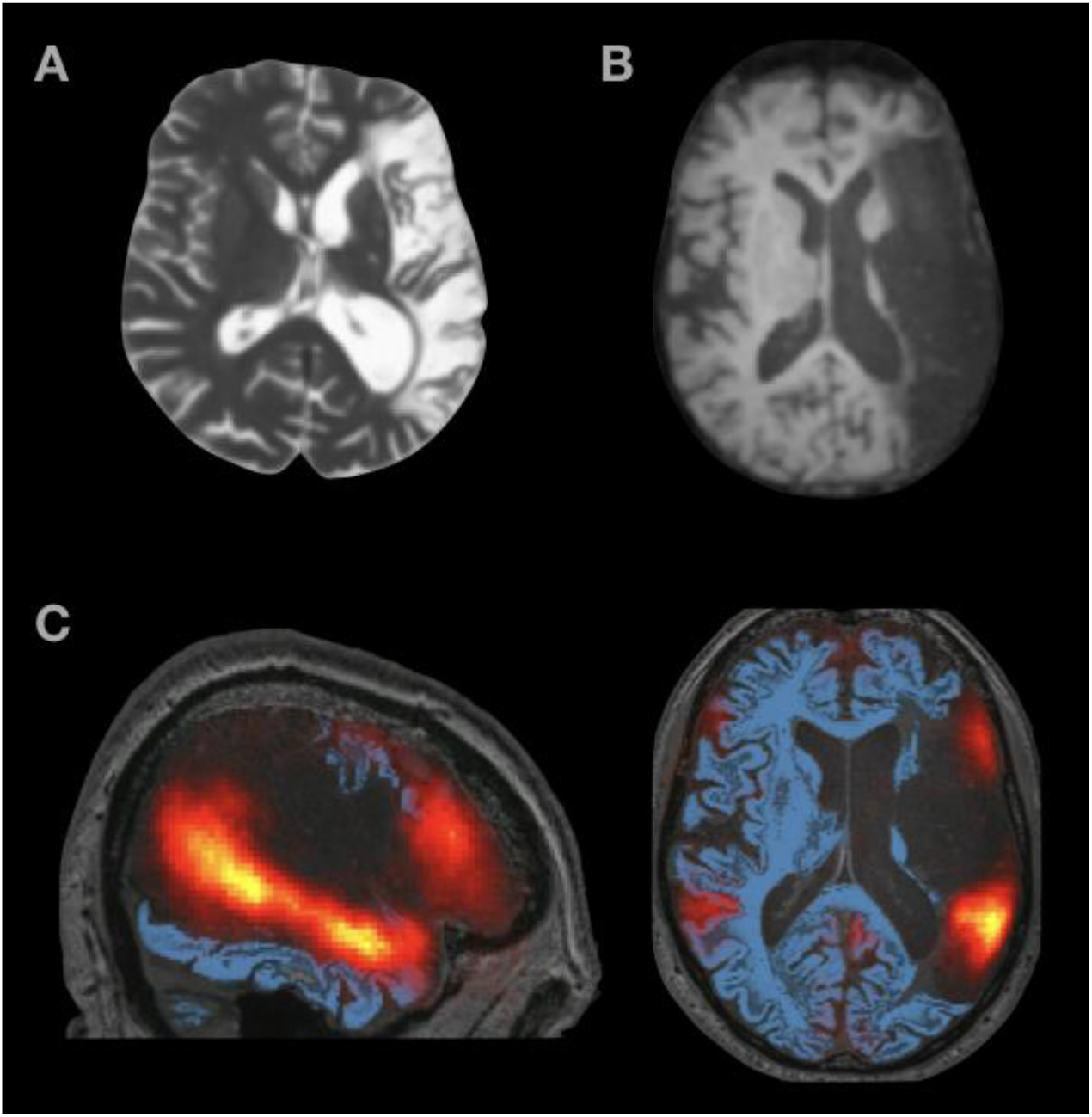
Structural MRI images from (A) S.A. and (B) P.R. (C) Probabilistic language activation overlap map overlaid on top of P.R.’s structural MRI image. The heatmap values range from 0.01 (red) to 0.5 (yellow). As can be seen, the lesion covers most left-hemisphere areas with voxels that likely belong to the language network.

Both individuals were severely agrammatic (**Table 2**). Whereas they had some residual lexical comprehension ability, scoring well on tasks involving word-picture matching and synonym matching across spoken and written modalities, their lexical production was impaired. Both failed to correctly name a single item in a spoken picture-naming task. S.A. displayed some residual written word production ability, scoring 24 out of 60 in a written picture-naming task. P.R., however, performed poorly in the written task, correctly naming just 2 out of 60 items.

**Table 2.** Results of linguistic assessments for participants with global aphasia.

| **Lexical Tests** | **Chance Score** | **S.A.** | **P.R.** |
| --- | --- | --- | --- |
| ADA spoken word picture matching | 16.5 | 60/66* | 61/66* |
| ADA written word picture matching | 16.5 | 62/66* | 66/66* |
| ADA spoken synonym matching | 80 | 123/160* | 113/160* |
| ADA written synonym matching | 80 | 121/160* | 145/160* |
| PALPA 54 spoken picture naming | n/a | 0/60 | 0/60 |
| PALPA 54 written picture naming | n/a | 24/60 | 2/60 |
| **Syntactic Tests** |  |  |  |
| Comprehension of spoken reversible sentences | 50 | 49/100 | 38/100 |
| Comprehension of written reversible sentences | 50 | 42/100 | 49/100 |
| Written grammaticality judgments | 20 | 26/40* | 21/40 |
| **Verbal Working Memory** |  |  |  |
| PALPA 13 digit span (recognition) | n/a | 3 items | 4 items |

\* indicates above chance performance ( $p < .05$ )
The tests were taken from the Action for Dysphasic Adults (ADA) Auditory Comprehension Battery (Franklin et al., 1992) and the Psycholinguistic Assessment of Language Processing in Aphasia (PALPA; Kay et al., 1992) or developed for the purpose of the study.

S.A. and P.R.’s syntactic processing was severely disrupted. They scored at or below chance in the reversible spoken and written sentence comprehension tasks (sentence-picture matching), which included active sentences, e.g. “the man kills the lion”, and passive sentences, e.g. “the man is killed by the lion”. They also scored near chance in written grammaticality judgment assessments. To determine whether the sentence comprehension impairments could be explained by phonological rather than syntactic deficits, we evaluated their phonological working memory by means of a digit span test (using a recognition paradigm that did not require language production). Both participants had a digit span that was sufficiently long to process the types of sentences used in the syntactic assessments, and thus their language difficulties could not be attributed to phonological working memory problems.

Importantly, S.A. and P.R. performed relatively well on nonverbal reasoning tasks, which included measures of fluid intelligence (Raven’s Standard/Colored Progressive Matrices; Raven & Raven, 2003), object semantics (Pyramids and Palm Trees test; Howard & Patterson, 1992), and visual working memory (Visual Pattern Test; Della Sala et al., 1999), indicating that the extensive brain damage in these patients did not ubiquitously affect all cognitive abilities (**Table 3**). Such a selective impairment of linguistic skills allowed us to examine the causal relationship between language and event semantics.

**Table 3.** Results of non-linguistic assessments for participants with global aphasia.

| **Reasoning Tests** | **S.A.** | **P.R.** |
| --- | --- | --- |
| Raven's Colored Progressive Matrices | 36/36 | 34/36 |
| Raven's Standard Progressive Matrices | 53/60 | 36/60 |
| Pyramids and Palm Trees (3 picture version) | 50/52 | 47/52 |
| Visual Pattern Test | 11.5 (90<sup>th</sup> percentile*) | 8.6 (40<sup>th</sup> percentile*) |

\* percentiles are calculated with respect to adults in the same age range with no neurological impairment

To test whether global aphasia affects general event semantics, we measured S.A. and P.R.’s performance on two tasks: (1) the picture plausibility task, identical to the pictures/semantic-task condition from Experiment 1, and (2) a sentence-picture matching task, during which participants saw a picture together with a sentence in which the agent and patient either matched the picture or were switched (“a cop is arresting a criminal” vs. “a criminal is arresting a cop”) and had to indicate whether or not the sentence matched the picture. The sentence-picture matching task was similar to the reversible sentence comprehension task in **Table 2**, with the exception that the pictures used were identical to pictures from the plausibility task and all sentences used active tense. For each task, patient performance was compared with the performance of 12 age-matched controls (58-78 years (mean 65.5 years) for the picture plausibility task; 58-78 years (mean 64.7 years) for the sentence-picture matching task).

The results showed a clear difference in performance between the picture plausibility task and the sentence-picture matching task (**Figure 4**). Both individuals with global aphasia and control participants performed well above chance when judging picture plausibility. Neurotypical controls had a mean accuracy of 95.7% (*SD* = 3.8%). Aphasia patients had mean accuracies of 91.0% (S.A.; 1.2 *SD* below average) and 84.6% (P.R.; 3.0 *SD* below average); exact binomial test showed that performance of both patients was above chance (S.A., *p* < .001, 95% CI [.82, .96]; P.R., *p* < .001, 95% CI [.75, .92]). Although their performance was slightly below the level of the controls, the data indicate that both patients were able to process complex semantic (agent-patient) relations to evaluate the plausibility of depicted events.

**Figure 4.**
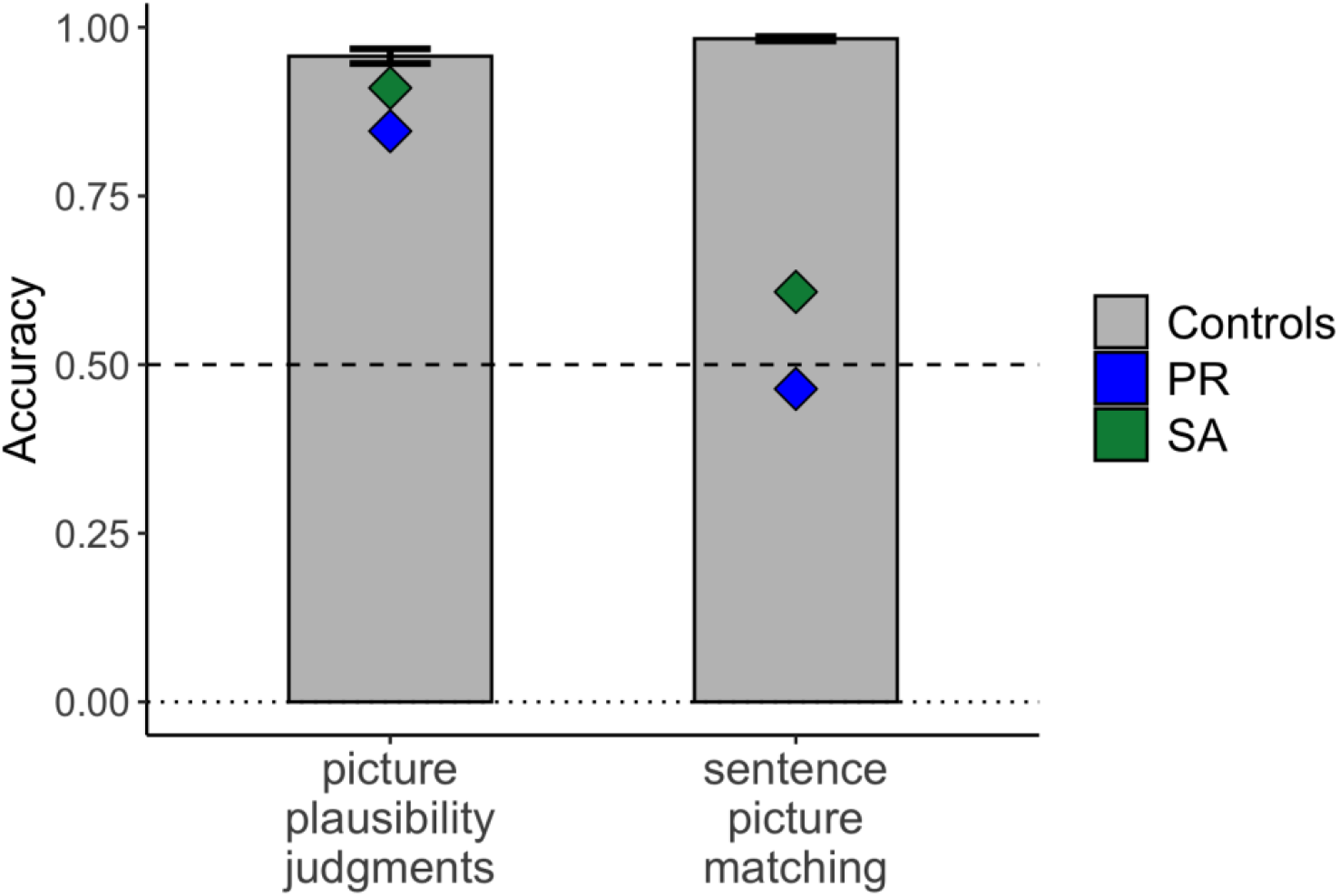
Individuals with profound aphasia perform well on picture plausibility judgment task but fail on the sentence-picture matching task. Patient accuracies are indicated in blue and green; control data (N=12) are shown in gray bars. The dotted line indicates chance performance. Error bars indicate standard error of the mean.

In the sentence-picture matching task, control participants performed close to ceiling, with a mean accuracy of 98.3% (SD = 1.1%). In contrast, both patients were severely impaired: S.A. had a mean accuracy of 60.8% and P.R. had a mean accuracy of 46.4%. Exact binomial test showed that P.R.’s performance was at chance (*p* = .464., 95% CI [.38, .55]), while S.A.’s performance was above chance (*p* = .009, 95% CI [.53, .69]) but still much lower than that of the controls. This result concurs with S.A.’s and P.R.’s poor performance on the reversible sentence comprehension tasks, which had a similar setup but used different materials. However, it stands in stark contrast with the participants’ ability to interpret agent-patient interactions in pictures. The Crawford-Howell (1998) *t*-test indicated a significant dissociation between the picture plausibility task and the sentence-picture matching task for both individuals (S.A., t(11) = 18.00, p < .001; P.R., t(11) = 24.20, p < .001). This dissociation held for both hit rate and false alarm rate (**Figure S2**).

The findings from Experiment 2 demonstrate that, in spite of severe linguistic impairments, individuals with global aphasia are able to access information about event participants depicted in a visual scene, the action taking place between them, the roles they perform in the context of this action, and the real-world plausibility of these roles, indicating that none of these processes require the presence of a functional language network.

## Discussion

The relationship between language and thought has been long debated, both in neuroscience (e.g., Binder & Desai, 2011; Bookheimer, 2002; Fedorenko & Varley, 2016; Friederici, 2020) and other fields (e.g., Carruthers, 2002; Hauser et al., 2002; Vygotski, 2012; Winograd, 1976). Here, we ask whether language-responsive regions of the brain are essential for a core component of thought: processing combinatorial semantic representations. We demonstrate that left-hemisphere language regions are active during semantic processing of events shown as pictures (although the semantic task elicits a stronger response in the context of sentences than in the context of pictures). We further show that the language network is not essential for event semantics, given that the two individuals with global aphasia, who lack most of their left-hemisphere language network, can still evaluate the plausibility of visually presented events. Our study advances the field in three ways: i) it explores relational semantic processing in the domain of events, moving beyond simple object semantics—the focus of most prior neuroimaging work on conceptual processing; ii) it evaluates neural overlap between linguistic and visual semantics at the level of individual participants; and iii) it provides causal evidence in support of a dissociation between language and non-verbal event semantics. In the remainder of the manuscript, we discuss the implications of these results.

### The language network is not essential for event semantics

Semantic processing of events is a complex, multi-component process. For instance, deciding whether or not an event is plausible requires one to (1) identify the relevant event participants, (2) determine the action taking place between them, (3) decipher the role that each event participant is performing (in our task, agent vs. patient), and finally, (4) estimate the likelihood that a given participant would be the agent/patient of the relevant action. Whereas the first three components can, at least in part, be attributed to input-specific processes (e.g., high-level vision), establishing plausibility cannot be solely attributed to perception: in order to decide whether a cop arresting a criminal is more likely than a criminal arresting a cop, participants need to draw on their world knowledge. We demonstrate that this highly abstract process can proceed even when the language network is severely impaired, thus providing strong evidence that semantics and language are neurally distinct.

The functional dissociation between language-based and vision-based semantic judgments of events accords with the fact that both non-human animals and preverbal infants are capable of complex event processing (Seed & Tomasello, 2010; Spelke, 1976) and that specialized neural mechanisms, distinct from the language network, have been associated with visual understanding of actions (Fang et al., 2016; Häberling et al., 2016; Tarhan & Konkle, 2020) and interactions between animate and/or inanimate entities (Fischer et al., 2016; Walbrin et al., 2018). These neural mechanisms are either bilateral or right-lateralized, which constitutes further evidence of their dissociation from language, which is typically left-lateralized.

If language is not essential for event semantics, why is the language network active during a non-verbal event semantic task? It is possible that neurotypical participants partially recode pictorial stimuli into verbal form (Greene & Fei-Fei, 2014; Trueswell & Papafragou, 2010), an operation that might provide access to linguistic representations as an additional source of task-relevant information (Connell & Lynott, 2013). Indeed, text-based computational models developed in recent years have been shown to successfully perform a wide range of “semantic” tasks, such as inference, paraphrasing, or question answering (Brown et al., 2020; Devlin et al., 2018, among others). Even simple n-gram models can determine the probability of certain events by, e.g., estimating the probability that the phrase “is arresting” directly follows “cop” vs. “criminal”. Given that semantic information derived from language is distinct from perception-based world knowledge (Lucy & Gauthier, 2017) and that people can flexibly use both depending on task demands (Willits et al., 2015), it is possible that the language network in neurotypicals provides an additional source of information when determining event plausibility. The absence of this additional information source may account for the small decrement in performance observed in our patients relative to the control participants.

That said, we consider it unlikely that semantic processing of picture stimuli in neurotypical participants was fully mediated by verbal representations. Verbal recoding of visual information is relatively slow and can only occur *after* semantic information has been retrieved from the picture (Potter et al., 1986; Potter & Faulconer, 1975). Furthermore, people do not typically generate internal verbal labels for visually presented objects without an explicit task to do so (Dahan et al., 2001; Magnuson et al., 2003; Rehrig et al., 2020; cf. Meyer et al., 2007), especially in the absence of memory demands (Pontillo et al., 2015). Our stimuli depicted complex two-participant events, making verbal recoding even more effortful than recoding of single objects and, therefore, unlikely to occur during a task that does not require linguistic label generation (Papafragou et al., 2008). Finally, even if the language network does play some role in nonverbal semantic processing in neurotypical participants, its recruitment is not essential, as indicated by the fact that patients with global aphasia can perform semantic evaluations of events (regardless of whether this evaluation proceeds through typical or compensatory pathways).

### Relationship to theories of semantics in the brain

In this paper, we specifically focused on the *role of the language network* in non-verbal event semantics, not on the question of which cognitive and neural mechanisms generally support event processing across representational formats (we investigate this question in other work; Ivanova et al., in prep.). Nonetheless, the results of this paper have important implications for general theories of semantic processing in the mind and brain.

Many current theories of semantics highlight broad anatomical areas implicated in linguistic processing as putative semantic hubs. Those include left angular gyrus (e.g., Binder & Desai, 2011), left inferior frontal cortex (e.g., Hagoort & van Berkum, 2007), and the anterior temporal lobes (e.g., Patterson et al., 2007). However, a large fraction of studies that provide evidence for these theories use linguistic stimuli and therefore cannot determine whether the same regions support non-linguistic semantic processing. Moreover, all the implicated areas are large patches of cortex that are structurally and functionally heterogeneous: as a result, simply because a visual-semantics study reports activation within the ‘LIFG’, or the ‘angular gyrus’, does not mean that the *language-responsive* portions of those broad areas are at play (see, e.g., Fedorenko & Blank, 2020 for discussion).

In the current study, the language-responsive fROIs we defined in left angular gyrus, left inferior frontal cortex, and left anterior temporal lobe all responded more strongly during the semantic task than during the perceptual task. This pattern is consistent with evidence of their general involvement in semantic processing but goes against some of the more specific claims made in the literature. For example, our results are inconsistent with the claim that the angular gyrus is the *primary* region involved in event semantics (Binder & Desai, 2011; cf. Williams et al., 2017) given that other regions show a similar functional response profile (although the fROI in the angular gyrus was the only one with numerically stronger responses to pictures than to sentences, so its involvement should be investigated further). Our results also go against the idea that the anterior temporal lobe is not engaged during the retrieval of event-level representations (Lewis et al., 2015; Schwartz et al., 2011; Teige et al., 2019; Xu et al., 2018).

In addition, our findings contribute to the body of work on the neural representation of agent/patient relationships. Previous experiments attempting to localize brain regions that support thematic role processing have attributed the processing of agent/patient relations to the left hemisphere. Frankland and Greene (2015, 2020) used sentence stimuli to isolate distinct areas in left superior temporal sulcus (STS) that are sensitive to the identity of the agent vs. the patient. Wang et al (2016) found that the same (or nearby) STS regions also contained information about thematic roles in videos depicting agent-patient interactions. However, the latter study identified a number of other regions that were sensitive to thematic role information, including clusters in the right posterior middle temporal gyrus and right angular gyrus, suggesting that left STS is not the only region implicated in thematic role processing. A similar distributed pattern was also reported in a neuropsychological study (Wu et al., 2007), which found that lesions to mid-STS led to difficulties in extracting thematic role information from both sentences and pictures; however, deficits in visual agent-patient processing were additionally associated with lesions in anterior superior temporal gyrus, supramarginal gyrus, and inferior frontal cortex, which casts further doubt on the unique role of left STS in agent-patient relation processing. We therefore conclude that parts of the left STS may play a role in processing linguistic information, including thematic relations and verb argument structure (Elli et al., 2019; Williams et al., 2017), but additional brain regions support the processing of thematic roles in non-verbal stimuli.

Finally, our results are generally consistent with a distributed view of semantic representations (McClelland & Rogers, 2003; Tyler & Moss, 2001). Multiple recent studies found that semantic information is not uniquely localized to any given brain region but rather distributed across the cortex (e.g., Anderson et al., 2017; Huth et al., 2016; Pereira et al., 2018). Distributing information across a network of regions in both left and right hemispheres enables the information to be preserved in case of brain damage (Schapiro et al., 2013), which would explain why patients with global aphasia preserve the ability to interpret visually presented events. That said, the findings reported here do not speak to the question of whether such representations mainly rely on sensorimotor areas (Barsalou, 2008; Pulvermuller, 1999) or on associative areas (Mahon, 2015; Mahon & Caramazza, 2008).

### Implications for neuroimaging studies of amodal semantics

The non-causal nature of the language network activation during a non-verbal semantic task has important implications for the study of amodal/multimodal concept representations. A significant body of work has aimed to isolate “amodal” representations of concepts by investigating the overlap between regions active during verbal and non-verbal presentations of a stimulus (Bright et al., 2004; Devereux et al., 2013; Fairhall & Caramazza, 2013; Handjaras et al., 2017; Sevostianov et al., 2002; Thierry & Price, 2006; Vandenberghe et al., 1996; Visser et al., 2012; A. D. Wagner et al., 1997). Most of these overlap-based studies have attributed semantic processing to frontal, temporal, and/or parietal regions within the left hemisphere. Our work, however, demonstrates that, even though meaningful linguistic and visual stimuli evoke overlapping activity in left-lateralized frontal and temporal regions, conceptual information about events persists even when most of these regions are damaged. Thus, overlapping areas of activation for verbal and non-verbal semantic tasks observed in brain imaging studies do not necessarily play a causal role in amodal event semantics.

Overall, our study emphasizes the importance of dissociating language and semantics when investigating the cognitive and neural mechanisms of thought. Our results show that semantic processing of visually presented events does not require the language network, drawing a sharp distinction between language and non-verbal semantics and highlighting the necessity to characterize the relationship between them in greater detail.

## Materials and Methods

### Experiment 1

#### Participants

Twenty-four participants took part in the fMRI experiment (11 female, mean age = 25 years, SD = 5.2). The participants were recruited from MIT and the surrounding Cambridge/Boston, MA, community and paid for their participation. All were native speakers of English, had normal hearing and vision, and no history of language impairment. All were right-handed (as assessed by Oldfield’s (1971) handedness questionnaire, or self-report). Two participants had low behavioral accuracy scores (<60%), and one had right-lateralized language regions (as evaluated by the language localizer task; see below); they were excluded from the analyses, which were therefore based on data from 21 participants. The protocol for the study was approved by MIT’s Committee on the Use of Humans as Experimental Subjects (COUHES). All participants gave written informed consent in accordance with protocol requirements.

#### Design, materials, and procedure

All participants completed a language localizer task aimed at identifying language-responsive brain regions (Fedorenko et al., 2010) and the critical picture/sentence plausibility task.

The localizer task was conducted in order to identify brain regions within individual participants that selectively respond to language stimuli. During the task, participants read sentences (e.g., NOBODY COULD HAVE PREDICTED THE EARTHQUAKE IN THIS PART OF THE COUNTRY) and lists of unconnected, pronounceable nonwords (e.g., U BIZBY ACWORRILY MIDARAL MAPE LAS POME U TRINT WEPS WIBRON PUZ) in a blocked design. Each stimulus consisted of twelve words/nonwords. For details of how the language materials were constructed, see Fedorenko et al (2010). The materials are available at http://web.mit.edu/evelina9/www/funcloc/funcloc_localizers.html. The sentences > nonword-lists contrast has been previously shown to reliably activate left-lateralized fronto-temporal language processing regions and to be robust to changes in the materials, task, and modality of presentation (Fedorenko et al., 2010; Mahowald & Fedorenko, 2016; Scott et al., 2017). Stimuli were presented in the center of the screen, one word/nonword at a time, at the rate of 450 ms per word/nonword. Each stimulus was preceded by a 100 ms blank screen and followed by a 400 ms screen showing a picture of a finger pressing a button, and a blank screen for another 100 ms, for a total trial duration of 6 s. Participants were asked to press a button whenever they saw the picture of a finger pressing a button. This task was included to help participants stay alert and awake. Condition order was counterbalanced across runs. Experimental blocks lasted 18 s (with 3 trials per block), and fixation blocks lasted 14 s. Each run (consisting of 5 fixation blocks and 16 experimental blocks) lasted 358 s. Each participant completed 2 runs.

The picture plausibility task included two types of stimuli: (1) black-and-white line drawings depicting plausible and implausible agent-patient interactions (created by an artist for this study), and (2) simple sentences describing the same interactions. Sample stimuli are shown in Figure 1, and a full list of materials is available on the paper website (https://osf.io/gsudr/). Forty plausible-implausible pairs of pictures, and forty plausible-implausible pairs of corresponding sentences were used. The full set of materials was divided into two lists, such that List 1 used plausible pictures and implausible sentences for odd-numbered items, and implausible pictures and plausible sentences for even-numbered items, and List 2 did the opposite. Thus, each list contained either a picture or a sentence version of any given event. Stimuli were presented in a blocked design (each block included either pictures or sentences) and were moving either to the right or to the left for the duration of stimulus presentation. At the beginning of each block, participants were told which task they would have to perform next: semantic or perceptual. The semantic task required them to indicate whether the depicted/described event is plausible or implausible by pressing one of two buttons. The perceptual task required them to indicate the direction of stimulus movement (right or left). To ensure that participants always perform the right task, a reminder about the task and the response buttons (“plausible=1/implausible=2”, or “moving right=1/left=2”) was visible in the lower right-hand corner of the screen for the duration of the block. Each stimulus (a picture or a sentence) was presented for 1.5 s, with 0.5 s intervals between stimuli. Each block began with a 2-second instruction screen to indicate the task, and consisted of 10 trials, for a total duration of 22 s. Trials were presented with a constraint that the same response (plausible/implausible in the semantic condition, or right/left in the perceptual condition) did not occur more than 3 times in a row. Each run consisted of 3 fixation blocks and 8 experimental blocks (2 per condition: semantic task – pictures, semantic task – sentences, perceptual task – pictures, perceptual task -sentences) and lasted 242 s (4 min 2 s). The order of conditions was palindromic and varied across runs and participants. Each participant completed 2 runs.

#### fMRI data acquisition

Structural and functional data were collected on the whole-body, 3 Tesla, Siemens Trio scanner with a 32-channel head coil, at the Athinoula A. Martinos Imaging Center at the McGovern Institute for Brain Research at MIT. T1-weighted structural images were collected in 176 sagittal slices with 1mm isotropic voxels (TR=2,530ms, TE=3.48ms). Functional, blood oxygenation level dependent (BOLD), data were acquired using an EPI sequence (with a 90° flip angle and using GRAPPA with an acceleration factor of 2), with the following acquisition parameters: thirty-one 4mm thick near-axial slices acquired in the interleaved order (with 10% distance factor), 2.1mm×2.1mm in-plane resolution, FoV in the phase encoding (A>>P) direction 200mm and matrix size 96mm×96mm, TR=2000ms and TE=30ms. The first 10s of each run were excluded to allow for steady state magnetization.

#### fMRI data preprocessing

MRI data were analyzed using SPM12 and custom MATLAB scripts (available in the form of an SPM toolbox from http://www.nitrc.org/projects/spm_ss). Each participant’s data were motion corrected and then normalized into a common brain space (the Montreal Neurological Institute (MNI) template) and resampled into 2mm isotropic voxels. The data were then smoothed with a 4mm FWHM Gaussian filter and high-pass filtered (at 200s). Effects were estimated using a General Linear Model (GLM) in which each experimental condition was modeled with a boxcar function (modeling entire blocks) convolved with the canonical hemodynamic response function (HRF).

#### Defining functional regions of interest (fROIs)

The critical analyses were restricted to individually defined language fROIs (functional regions of interest). These fROIs were defined using the Group-constrained Subject-Specific (GSS) approach (Fedorenko et al., 2010; Julian et al., 2012), where a set of spatial parcels is combined with each individual subject’s localizer activation map to constrain the definition of individual fROIs. The parcels mark the expected gross locations of activations for a given contrast based on prior work and are sufficiently large to encompass the extent of variability in the locations of individual activations. Here, we used a set of six parcels derived from a group-level probabilistic activation overlap map for the sentences > nonwords contrast in 220 participants. These parcels included two regions in the left inferior frontal gyrus (LIFG, LIFGorb), one in the left middle frontal gyrus (LMFG), two in the left temporal lobe (LAntTemp and LPostTemp), and one extending into the angular gyrus (LAngG). The parcels are available on the paper’s website. Within each parcel, we selected the top 10% most responsive voxels, based on the *t* values for the sentences > nonwords contrast (see Figure 1 in Blank et al (2014) or Figure 1 in Mahowald & Fedorenko (2016), for sample fROIs). Individual-level fROIs defined in this way were then used for subsequent analyses that examined the behavior of the language network during the critical picture/sentence plausibility task.

#### Examining the functional response profiles of the language fROIs

For each language fROI in each participant, we averaged the responses across voxels to get a value for each of the four critical task conditions (semantic task on pictures, semantic task on sentences, perceptual task on pictures, perceptual task on sentences). We then ran a linear mixed-effect regression model with two fixed effects (stimulus type and task) and two random effects (participant and fROI). Planned follow-up comparisons examined response to sentences and pictures during the semantic task within each fROI; the results were FDR-corrected (Benjamini & Hochberg, 1995) for the number of regions. The formula used for the main mixed linear effects model was *EffectSize ∼ StimType*Task + (1*|*fROI) + (1*|*Subject)*. The formula used for the follow-up comparisons was *EffectSize ∼ StimType*Task + (1*|*Subject)*. The analysis was run using the *lmer* function from the *lme4* R package (Bates et al., 2015); statistical significance of the effects was evaluated using the *lmerTest* package (Kuznetsova et al., 2017).

### Experiment 2

#### Participants

Two participants with global aphasia, S.A. and P.R., took part in the study. Both had large lesions that had damaged the left inferior frontal gyrus, the inferior parietal lobe (supramarginal and angular gyri) and the superior temporal lobe. At the time of testing, they were 68 and 70 years old respectively. S.A. was 22 years 5 months post-onset of his neurological condition, and P.R. was 14 years 7 months post-onset. S.A. had a subdural empyema in the left sylvian fissure, with associated meningitis that led to a secondary vascular lesion in left middle cerebral artery territory. P.R. also had a vascular lesion in left middle cerebral artery territory.

Both participants were male, native English speakers, and did not present with visual impairments. S.A. was pre-morbidly right-handed; P.R. was pre-morbidly left-handed, but a left hemisphere lesion that resulted in profound aphasia indicated that he, like most left-handers, was left-hemisphere dominant for language (Pujol et al., 1999). Both individuals were classified as severely agrammatic (**Table 1**), but their non-linguistic cognitive skills were mostly spared (**Table 2**). They performed the semantic task and the sentence-picture matching task with a 7-months period between the two.

We also tested two sets of neurotypical control participants, one for the semantic task and one for the language task. The semantic task control participants were 12 healthy participants (7 females) ranging in age from 58 to 78 years (mean age 65.5 years). The language task control participants were 12 healthy participants (5 females) ranging in age from 58 to 78 years (mean age 64.7 years). None of the healthy participants had a history of speech or language disorders, neurological diseases or reading impairments. All were native English speakers, and had normal, or corrected-to-normal, vision.

Participants undertook the experiments individually, in a quiet room. An experimenter was present throughout the testing session. The stimuli were presented on an Acer Extensa 5630G laptop, with the experiment built using DMDX (Forster & Forster, 2003). Ethics approval was granted by the UCL Research Ethics Committee (LC/2013/05). All participants provided informed consent prior to taking part in the study.

#### Semantic Task: Picture plausibility judgments

The same picture stimuli were used as those in Experiment 1 (see **Figure 1**), plus one additional plausible-implausible pair of pictures (which was omitted from the fMRI experiment to have a total number of stimuli be divisible by four, for the purposes of grouping materials into blocks and runs), for a total of 82 pictures (41 plausible-implausible pairs). Four of the 82 pictures were used as training items (see below).

The stimuli were divided into 2 sets, with an equal number of plausible and implausible pictures; each plausible-implausible pair was split across the 2 sets, to minimize repetition of the same event participants within a set. The order of the trials was randomized within each set, so that each participant saw the pictures in a different sequence. A self-timed break was placed between the two sets.

Prior to the experiment, participants were shown two pairs of pictures, which acted as training items. The pairs consisted of one plausible and one implausible event. They were given clear instructions to focus on the relationship between the two characters and assess whether they thought the interaction was plausible, in adherence with normal expectations, or implausible, at odds with expectations. They were asked to press a green tick (the left button on the mouse) if they thought the picture depicted a plausible event, and a red cross (the right button on the mouse) if they thought the picture depicted an implausible event. They were asked to do so as quickly and accurately as possible. The pictures appeared for a maximum of 8 seconds, with the inter-stimulus interval of 2 seconds. Accuracies and reaction times were recorded. Participants had the opportunity to ask any questions, and the instructions for participants with aphasia were supplemented by gestures to aid comprehension of the task. Participants had to indicate that they understood the task prior to starting.

#### Language Task: Sentence-picture matching

The same 82 pictures were used as in the plausibility judgment experiment. In this control task, a sentence was presented below each picture that either described the picture correctly (e.g., “the policeman is arresting a criminal” for the first sample picture in **Figure 1**) or had the agent and patient switched (“the criminal is arresting the policeman”). Simple active subject-verb-object sentences were used. Combining each picture with a matching and a mismatching sentence resulted in 164 trials in total.

For the control participants, the trials were split into two sets of 82, with an equal number of plausible and implausible pictures, as well as an equal number of matches and mismatches in each set. In order to avoid tiring the participants with aphasia, the experiment was administered across two testing sessions each consisting of two sets of 41 stimuli and occurring within the same week. For both groups, the order of the trials was randomized separately for each participant, and no pictures from the same pair (e.g., an event involving a cop and a criminal) appeared in a row. A self-timed break was placed between the two sets.

Prior to the experiment, participants were told that they would see a series of pictures with accompanying sentences, and their task was to decide whether the sentence matched the depicted event. They were asked to press a green tick (the left button on the mouse) if they thought the sentence matched the picture, and a red cross (the right button on the mouse) if they thought the sentence did not match the picture. They were asked to do so as quickly and accurately as possible. The picture/sentence combinations appeared for a maximum of 25 seconds, with the inter-stimulus interval of 2 seconds. Accuracies and reaction times were recorded. As in the critical task, participants had the opportunity to ask any questions, and the instructions for participants with aphasia were supplemented by gestures.

#### Data analysis

We used exact binomial test to test whether patients’ performance on either task was significantly above chance, as well as the Crawford-Howell (1998) test for dissociation to compare patient performance relative to controls across the two tasks. We excluded all items with reaction times and/or accuracies outside 3 standard deviations of the control group mean (4 items for the semantic task and 11 items for the sentence-picture matching task).

#### Estimating the damage to the language network in patients with aphasia

In order to visualize the extent of the damage to the language network, we combined available structural MRI of one patient with aphasia with a probabilistic activation overlap map of the language network. The map was created by overlaying thresholded individual activation maps for the language localizer contrast (sentences > nonwords, as described in Experiment 1) in 220 healthy participants. The maps were thresholded at the p<0.001 whole-brain uncorrected level, binarized, and overlaid in the common space, so that each voxel contains information on the proportion of participants showing a significant language localizer effect (see Woolgar, Duncan, Manes & Fedorenko (2018) for more details). The map can be downloaded from the paper’s website. An overlay of this probabilistic map onto P.R.’s structural scan is shown in **Figure 3**.

## Acknowledgments

We would like to acknowledge the Athinoula A. Martinos Imaging Center at the McGovern Institute for Brain Research at MIT, and its support team (Steve Shannon and Atsushi Takahashi). We thank Birgit Zimmerer for creating the picture stimuli used in both experiments, Chloe Bustin for norming the stimuli, Lily Jordan for help with the behavioral piloting of the fMRI experiment, EvLab members for their help with fMRI data collection. E.F. was supported by NIH awards R00-HD057522, R01-DC016607, and R01-DC016950, by a grant from the Simons Foundation to the Simons Center for the Social Brain at MIT, and by funds from BCS and the McGovern Institute for Brain Research at MIT. R.V. was supported by Arts and Humanities Research Council and Alzheimer’s Society awards.

## Supplemental Information

### Experiment 1

**Figure S1.**
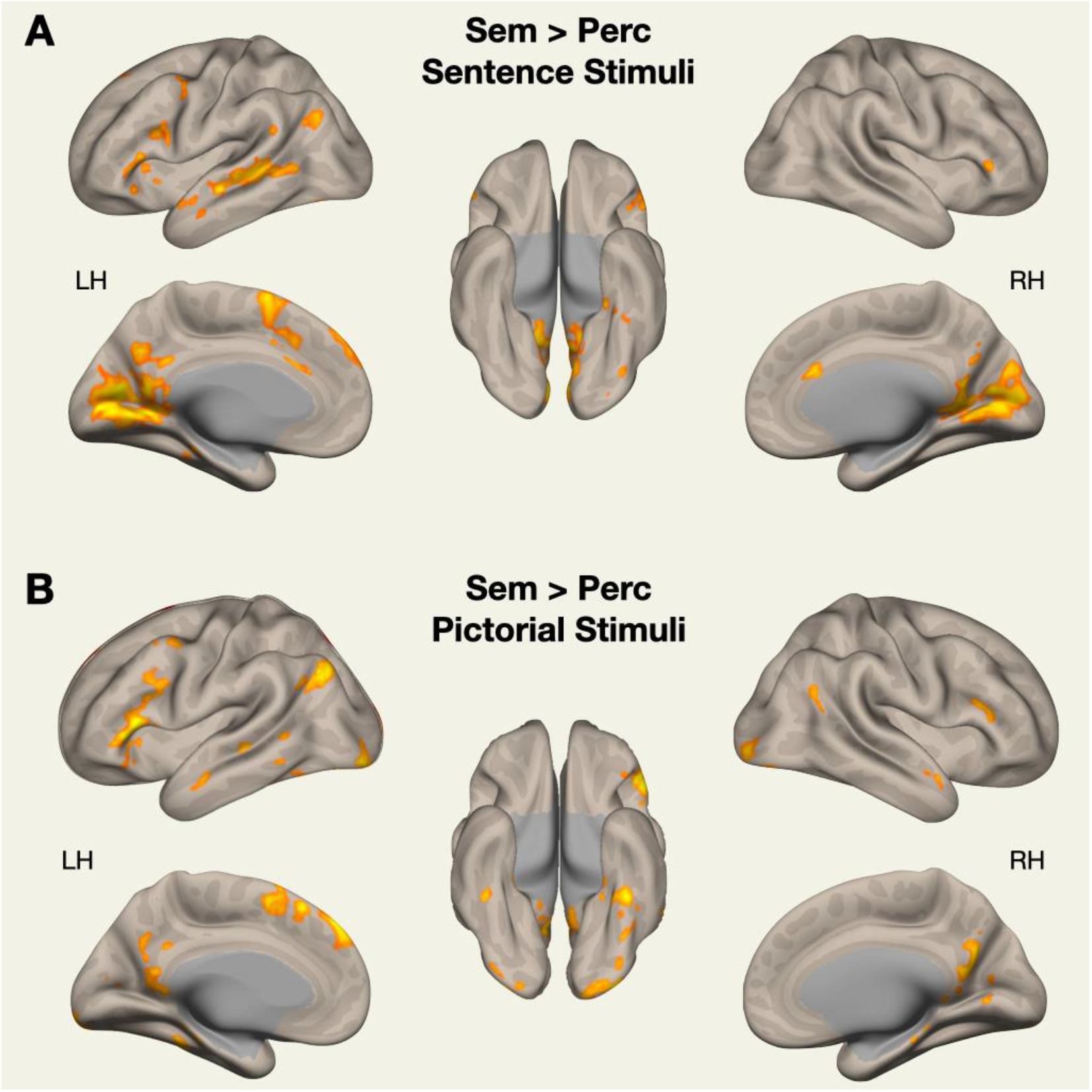
Whole-brain random effects group analysis (Holmes&Friston, 1998) for Semantic > Perceptual task contrast, conducted separately on the data from sentence trials (a) and picture trials (b). The analysis was conducted using the spm_ss toolbox (available at http://www.nitrc.org/projects/spm_ss), which interfaces with SPM and the CONN toolbox (https://www.nitrc.org/projects/conn). The results were thresholded at p=0.001, and resulting clusters were FDR-corrected at p=0.05.

#### Behavioral Results

Average response rate was 92% and did not vary significantly across tasks. Average response times were 1.27 s (SD = 0.46) for the semantic sentence task, 1.16 s (SD = 0.38) for the perceptual sentence task, 1.22 s (SD = 0.35) for the semantic picture task, and 1.19 (SD = 0.36) for the perceptual picture task. Average accuracies were 0.81 for the semantic sentence task, 0.79 for the perceptual sentence task, 0.75 for the semantic picture task, and 0.75 for the perceptual picture task. Due to a technical error, behavioral data from 14 out of 21 participants was only recorded for one of the two runs.

### Experiment 2

**Figure S2.**
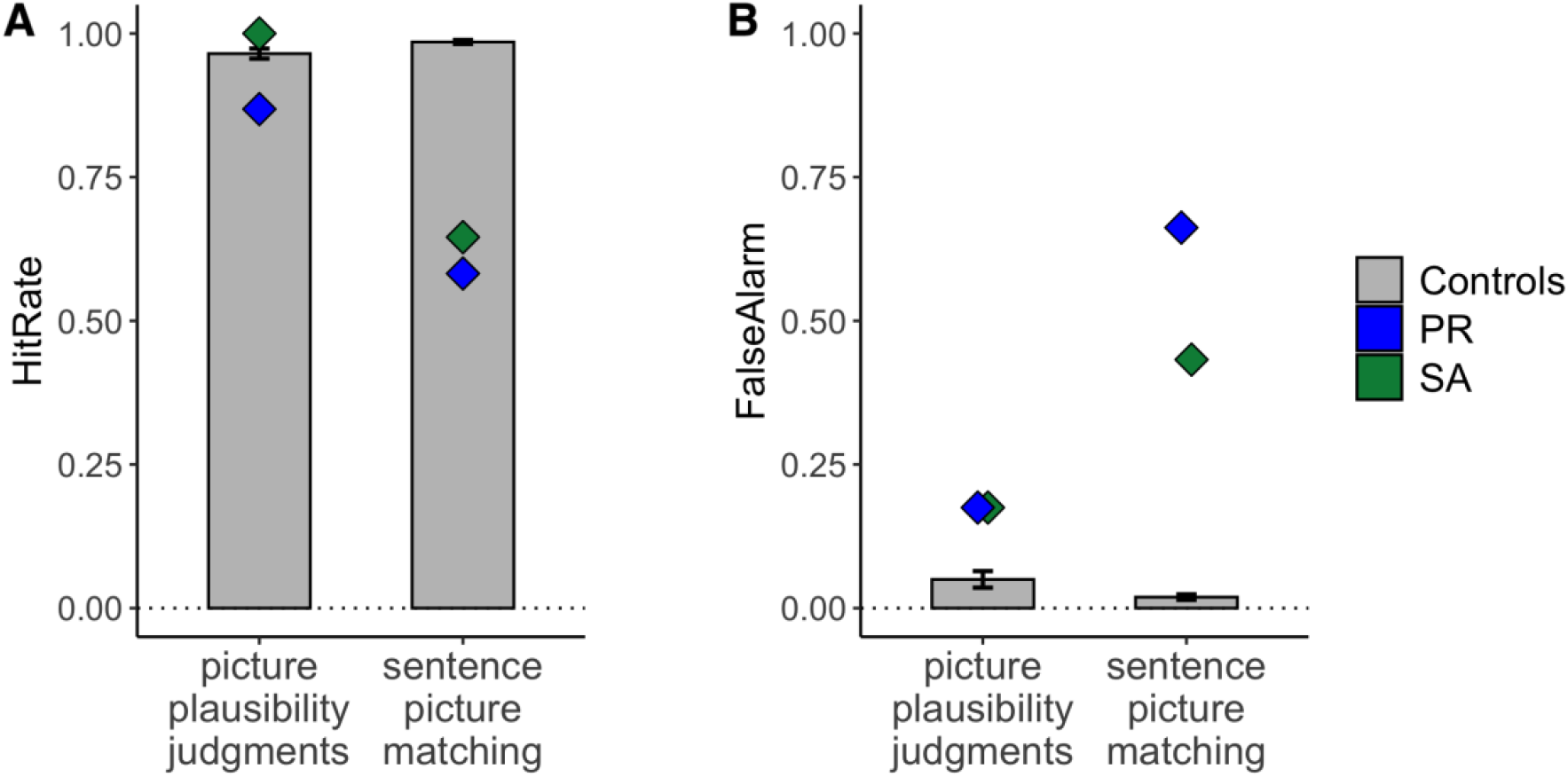
Hit rate (A) and false alarm rate (B) for Experiment 2 tests. Error bars indicate standard error of the mean. The Crawford-Howell test indicated a significant dissociation between the two tests for both hit rate (S.A.: t(11) = 18.95, p < .001; P.R.: t(11) = 19.59, p < .001) and false alarm rate (S.A.: t(11) = 12.55, p < .001; P.R.: t(11) = 20.31, p < .001).

